# RNA structure modulates Cas13 activity and enables mismatch detection

**DOI:** 10.1101/2023.10.05.560533

**Authors:** Ofer Kimchi, Benjamin B. Larsen, Owen R. S. Dunkley, Aartjan J.W. te Velthuis, Cameron Myhrvold

**Affiliations:** Lewis-Sigler Institute for Integrative Genomics, Princeton University, Princeton, New Jersey, 08544, USA; Department of Molecular Biology, Princeton University, Princeton, New Jersey, 08544, USA; Department of Chemical and Biological Engineering, Princeton University, Princeton, New Jersey, 08544, USA; Omenn-Darling Bioengineering Institute, Princeton University, Princeton, New Jersey, 08544, USA; Department of Chemistry, Princeton University, Princeton, New Jersey, 08544, USA

**Keywords:** Cas13, CRISPR, RNA detection, RNA structure, strand displacement

## Abstract

The RNA-targeting CRISPR nuclease Cas13 has emerged as a powerful tool for applications ranging from nucleic acid detection to transcriptome engineering and RNA imaging^1–6^. Cas13 is activated by the hybridization of a CRISPR RNA (crRNA) to a complementary single-stranded RNA (ssRNA) protospacer in a target RNA^1,7^. Though Cas13 is not activated by double-stranded RNA (dsRNA) *in vitro*, it paradoxically demonstrates robust RNA targeting in environments where the vast majority of RNAs are highly structured^2,8^. Understanding Cas13’s mechanism of binding and activation will be key to improving its ability to detect and perturb RNA; however, the mechanism by which Cas13 binds structured RNAs remains unknown^9^. Here, we systematically probe the mechanism of LwaCas13a activation in response to RNA structure perturbations using a massively multiplexed screen. We find that there are two distinct sequence-independent modes by which secondary structure affects Cas13 activity: structure in the protospacer region competes with the crRNA and can be disrupted via a strand-displacement mechanism, while structure in the region 3’ to the protospacer has an allosteric inhibitory effect. We leverage the kinetic nature of the strand displacement process to improve Cas13-based RNA detection, enhancing mismatch discrimination by up to 50-fold and enabling sequence-agnostic mutation identification at low (<1%) allele frequencies. Our work sets a new standard for CRISPR-based nucleic acid detection and will enable intelligent and secondary-structure-guided target selection while also expanding the range of RNAs available for targeting with Cas13.

## Main text

The RNA-targeting CRISPR effector protein Cas13 holds tremendous promise for numerous applications, such as RNA targeting, detection, editing, and imaging^1–5,7,10^. Cas13 is activated by the hybridization of a CRISPR RNA (crRNA) spacer sequence to a complementary region in a target RNA (the protospacer)^7,11^. Once activated, Cas13 cleaves both the target RNA (*cis* cleavage) and other RNAs in solution (*trans* cleavage)^2,3,7,9^. However, the biophysical process by which the Cas13-crRNA complex binds to and is activated by a target RNA is poorly understood.

Understanding the mechanism of Cas13 activation would solve several key challenges. First, even with perfect complementarity between crRNA and protospacer, Cas13 activity levels can vary by several orders of magnitude^12–14^. While the specifics of the crRNA and target sequences—including spacer length, nucleotide sequence, and crRNA/protospacer mismatches—are known to affect Cas13 activity, much of this variation remains unaccounted for^12–14^. RNA secondary structure is an intriguing candidate to explain this variation, since unlike the CRISPR effector proteins Cas9 and Cas12, Cas13 is believed to be incapable of unwinding structured targets^11,15^. Models including only sequence effects while excluding structural effects have had success with a binary classification of active/inactive crRNAs, but have not been able to solve the regression problem of quantitatively predicting Cas13 activity from the crRNA sequence^12,13,16^. Second, although Cas13 activation requires crRNA/protospacer complementarity, Cas13 is frequently activated to a similar or even greater degree in the presence of single mismatches^17,18^. We hypothesized that these two poorly understood characteristics of Cas13 could be addressed by studying the activation of Cas13 in more detail using structured RNAs as a model system.

RNA molecules form intramolecular base pairs (secondary structures) that compete with intermolecular RNA-RNA interactions. We sought to explore how this competition affects the crRNA-target interactions underlying Cas13 activation. RNA secondary structure has long been suspected to influence Cas13 activity due to its competition with the crRNA for target base pairing, but this has been challenging to study in isolation, as primary sequence and secondary structure are inextricably linked^19^. Abudayyeh & Gootenberg *et al.* showed a small but significant negative correlation between secondary structure and Cas13a activity for crRNAs targeting different regions of four genes in HEK293FT cells^2^. By tiling the long non-coding RNA Xist in HEK293T cells, Bandaru et al. showed that crRNAs are typically more active when targeting single-stranded regions than double-stranded regions^20^. By examining different regions of natural RNAs, prior studies have been unable to distinguish between effects on Cas13 caused by changes to target structure, and those caused by changes to target sequence. Thus, the degree to which the RNA structure affects crRNA-target binding and Cas13 activity remains unclear.

### RNA structure reduces LwaCas13a activity

To isolate the effect of RNA structure on Cas13 activity, we designed an ssRNA protospacer sequence to which we could add variable amounts of secondary structure by either intramolecular extension of an RNA hairpin, or by adding external complementary RNA or DNA oligonucleotides of different lengths, termed “occluders” (Fig. 1A). We designed the protospacer to reflect viral sequence diversity and to have minimal secondary structure in the absence of these modifications (see Methods). We tested the ability of these structured protospacers to activate LwaCas13a using cleavage of a quenched fluorescent RNA to report activity. Increased secondary structure decreased Cas13 activity across all three assay conditions (Fig. 1A). We next quantified Cas13 activity by fitting the fluorescence curves to effectively first-order reaction equations, defining activity as the rate of reporter cleavage (hr^-1^), a proxy for the concentration of active Cas13 in the system (see Methods). We observed a high degree of correlation between the three types of target occlusion (Fig. 1B). Cas13 activity varied by an order of magnitude for the same sequence with different amounts of target occlusion (Fig. 1C).

**Fig. 1:**
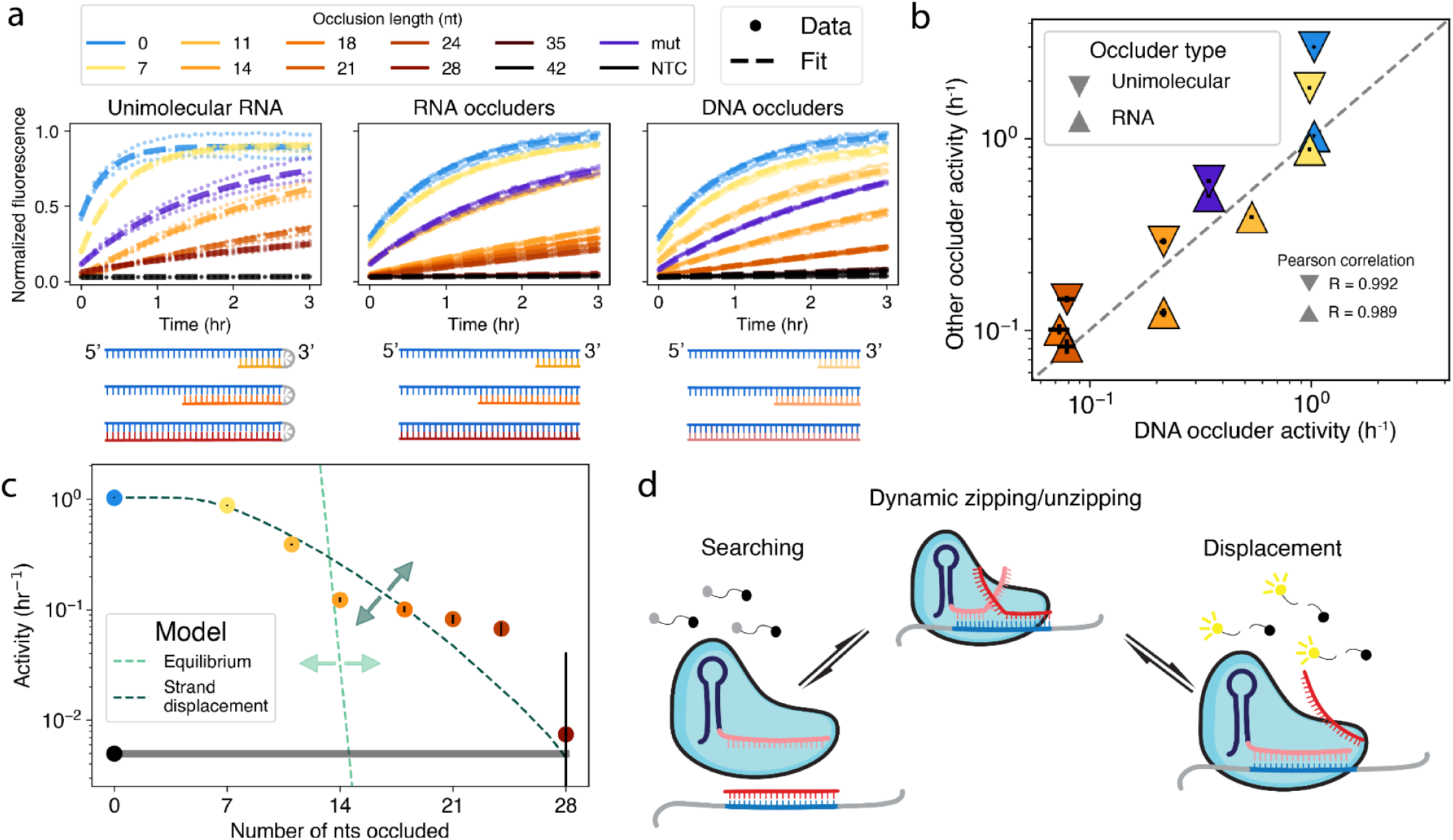
RNA structure leads to a reduction in Cas13 activity due to strand displacement. **A:** Different amounts of structure were introduced into the target via intramolecular structure, RNA oligos, and DNA oligos; resulting fluorescent kinetic curves. Target input concentration: 7.5 x 10^8^ copies/μL. “Mut” denotes 28nt of occlusion with nine mismatches to weaken binding. Data from n=3 technical replicates are shown (dots). **B:** Scatter plot comparing the impact of the different occlusion types depicted in (A) on Cas13 activity; x axis: Cas13 activity when occluded by DNA oligos, y axis: Cas13 activity when occluded by intramolecular RNA or RNA oligos. **C:** Cas13 activity vs. occluder length for RNA occluders is compared to two single-parameter models: an equilibrium model based on crRNA-target hybridization free energies(pink) and a free-energy-independent strand displacement model (green). Effects of changing the single parameter are indicated by arrows. Gray bar is NTC. **D:** Overview of strand displacement reactions. After initial binding to part of the target (blue), the crRNA (pink) and occluder (red) undergo a random walk process until one or the other is fully displaced. Displacement of the occluder leads to Cas13 activation. Error bars in B and C show standard deviation.

### Activity reduction is quantitatively explained by a kinetic strand displacement model

An equilibrium model based on the free energy of each target RNA (see Methods) failed to quantitatively account for the degree of structure-mediated Cas13 activity reduction (Fig. 1C light green dotted line). Labeling the free energy of the target-occluder complex by 𝛥𝐺_u_, the disagreement between the large difference in thermodynamic drives (𝑒𝑥𝑝 (-β𝛥𝐺_u_) ranges over 30 orders of magnitude) and the smaller difference in activities (ranging over 2 orders of magnitude) cannot be explained by an equilibrium RNA-RNA hybridization framework. Given known free energies of RNA-RNA binding^21^, the system temperature would have to be ∼7500K to match an equilibrium model to the measured Cas13 activity reduction.

We next sought to find a suitable alternative framework to explain the measured activity levels; a strand displacement model presents one such framework^22^. In this model, after initial binding to the target, the crRNA and occluding strand compete through a random-walk-like process until either the occluding strand is fully displaced or the crRNA-Cas13 complex dissociates from the target RNA^23,24^ (Fig. 1D). We hypothesized that strand displacement must occur for Cas13 to bind structured RNA.

In our model, Cas13 first binds non-specifically to a region of RNA^15^ and then performs a local search for a sequence complementary to its bound crRNA. If Cas13 is not activated within a given time t_dwell_ (i.e. it does not fully bind to a protospacer sequence complementary to its crRNA) it dissociates from the RNA and repeats the search process. This sequence of events is analogous to the process by which enzymes such as the *E. coli lac* repressor search for their binding site on DNA^25,26^. However, the strand displacement model predicts that as secondary structure length increases, the probability of Cas13 completing the strand displacement reaction within the time t_dwell_ decreases, leading to a proportional decrease in Cas13 activity. This model fits our measured results with a value of t_dwell_ equivalent to 100 steps of a strand displacement reaction (∼2 x 10^-5^ 𝑠), in good agreement with direct measurements of typical dwell times for DNA-binding proteins^25^ (Fig. 1C dark green dotted line; see Methods).

### A massively multiplexed assay reveals structure effects are sequence independent

To probe the limits of the strand displacement model, we used a massively multiplexed assay to explore a broad range of target structure conditions for multiple sequences. Our assay uses DNA oligos to create secondary structure at defined positions in the target, having previously validated their effect as a proxy for RNA structure (Fig. 1B, 2A). We designed a single 1kb-long RNA molecule with minimal internal secondary structure^27^ (Extended Data Fig. 1A-C); a NUPACK prediction estimates its minimum free energy due to intramolecular contacts to be ∼-6 kcal/mol, on par with random 35-nt-long RNA sequences^28^. Our target RNA is divided into two control blocks (one at each end of the molecule) and eight experimental blocks, allowing for efficient multiplexing^12^. Each block contains a 28-nt-long protospacer flanked by two 34-nt buffer regions. For occlusion, we used DNA oligos of lengths 10, 14, 21, and 28 nucleotides (nt) in 3-nt-spaced tilings, for a total of 4,608 simultaneous conditions. We tested these conditions in parallel using a microfluidic chip-based assay^6^. Summaries of the resulting dataset are shown in Fig. 2B and Extended Data Figs. 2 and 3.

**Fig. 2:**
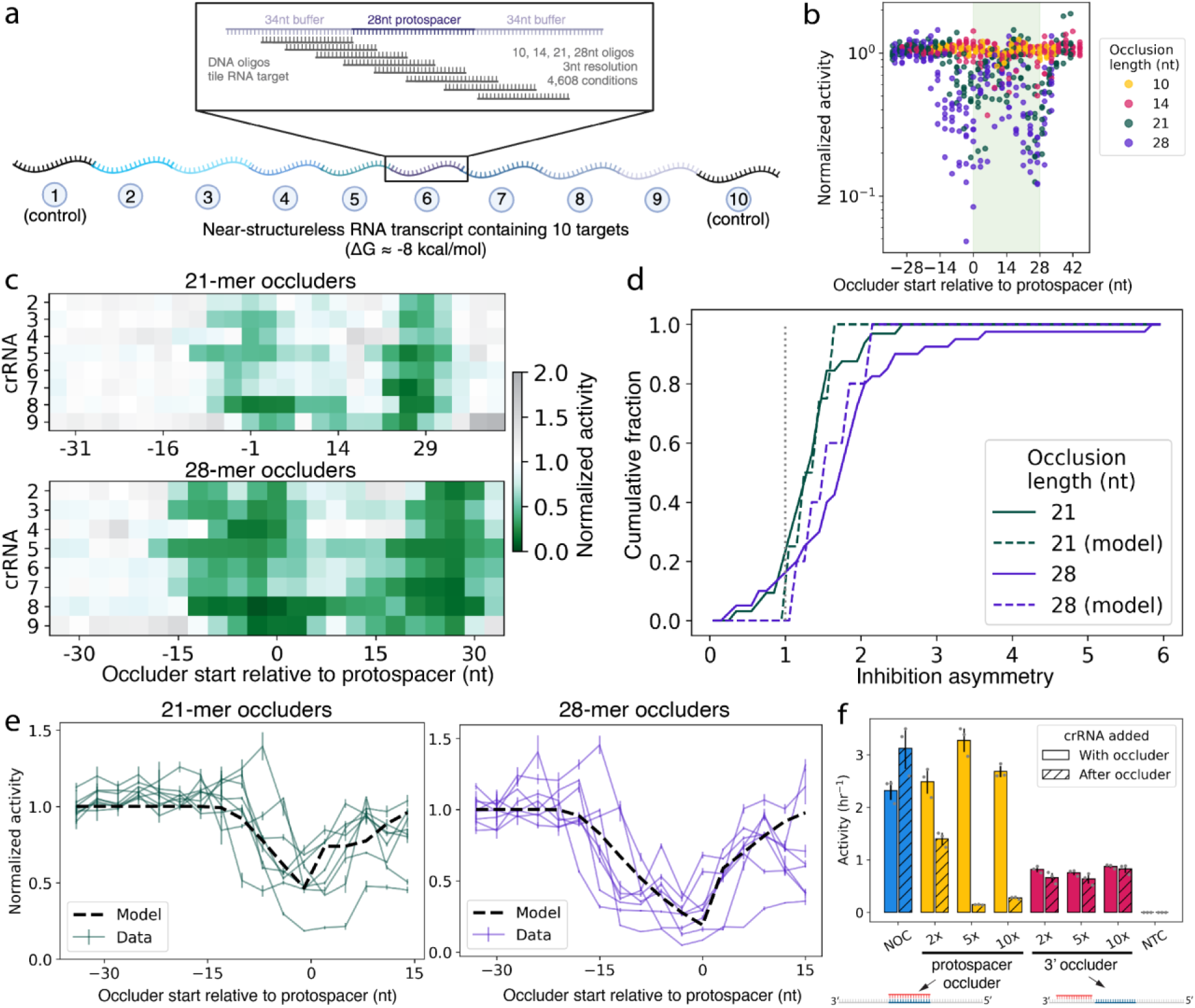
A massively multiplexed assay modulating secondary structure. **A:** Overview of the multiplexed assay, in which a total of 4,608 simultaneous assays were performed, with oligo occluders of lengths 10, 14, 21, 28 tiling each protospacer region in 3-nt increments. Made using BioRender. **B:** Overview of the Cas13 activity data from the multiplexed assay. Each data point represents the mean activity resulting from averaging four time series curves (see Methods), normalized to the non-occluded condition; positive and negative controls are not shown (Extended Data Fig 3). **C:** Heat map showing the degree of activity reduction (darker greens) by each 21mer and 28mer occluder. **D:** Cumulative histogram of inhibition asymmetry, defined as the ratio of activities when the same numbers of nucleotides are occluded at the 3’ vs. 5’ ends of the protospacer (see Extended Data Fig. 4C,D). **E:** Normalized Cas13 activity for 21mer (green) and 28mer (purple) occluders with different start positions in the region around the protospacer. Each line represents one crRNA. **F:** Bar chart showing the inhibitory effect of 28mer occluders overlapping the protospacer or the region 3’ to the protospacer, at different occluder concentrations and when annealing the occluder before or at the same time as the crRNA. In E and F, error bars show standard deviation.

Our results demonstrate that the reduction in Cas13 activity as a result of target structure is relatively sequence-independent, with the 8 experimental target blocks showing similar activity profiles in spite of large variation in absolute activity across these blocks (Fig. 2C, Extended Data Fig. 2). While 10- and 14-nt-long occluders had negligible effects on Cas13 activity, 21- and 28-mers had a strong effect. Consistent with earlier results, occluders binding to more of the protospacer typically led to a greater activity reduction. In agreement with our dwell time model and in contrast to other strand displacement systems^24,29,30^, the presence or absence of toeholds (unoccluded RNA) had little effect on Cas13 activity (Extended Data Figs. 4E-F).

The data also revealed an unexpected asymmetry among the effect of occluders on Cas13, in which occluders binding to the 5’ end of the protospacer had a larger effect on Cas13 activity than occluders binding the same number of nucleotides at the 3’ end (Fig. 2D, Extended Data Fig. 4A-D). We accounted for the asymmetry in our model by adding a second parameter to the model described above, creating a differential in t_dwell_ depending on whether or not the 5’ end of the protospacer is occluded. We found that our revised dwell time model was able to quantitatively capture the effects of secondary structure on Cas13 activity (Fig. 2D, E).

### Structure occluding the region 3’ of the protospacer inhibits Cas13

Surprisingly, when occluders are placed directly 3’ to the protospacer, Cas13 activity is potently inhibited. This second regime of inhibition exists across all tested crRNAs, and inhibition is strong for both 21mer and 28mer occluders (Fig. 2C). The non-monotonicity of this second activity trough cannot be explained using a strand displacement model, implying this drop in activity is not due to a reduction in crRNA-target binding.

We sought to probe whether this effect on Cas13 activity is a result of competitive or allosteric inhibition. For the protospacer occluder, but not for the 3’ occluder, we observe full rescue of Cas13 activity when the crRNA and occluder are added at the same time. Increasing the concentrations of 3’ occluder does not increase their inhibitory effect (Fig. 2F). These results indicate that the activity reduction conferred by occluding the region 3’ to the protospacer is likely the result of an allosteric effect.

### Strand displacement enhances mismatch detection

We hypothesized that the insights from our strand displacement model could help us dramatically improve the specificity of Cas13-based RNA detection assays. Past work has shown that secondary structure can make nucleic acid hybridization more sensitive to mismatches, both in CRISPR-based approaches and in other assays^31–34^; we hypothesized that given the kinetic nature of our assays, we could take advantage of the kinetic nature of strand displacement to similar ends without the necessity of a binding toehold required in other approaches. With no internal structure, even a mismatched crRNA is expected to bind strongly to the target; however, an occluding strand provides an extra kinetic barrier that is less likely to be overcome by a mismatched crRNA than by one that is perfectly complementary, thus improving specificity given the short dwell time of inactive Cas13 on the RNA (Fig. 3A). A differential-equation-based strand displacement model inspired by Ref.^35^ supported this hypothesis, revealing that even when both perfectly-complementary and mismatched invader strands bind strongly to the target in equilibrium, in the presence of an occluding oligo, the mismatched invader takes much longer to do so than the perfectly-matched invader (Fig. 3B).

**Fig. 3:**
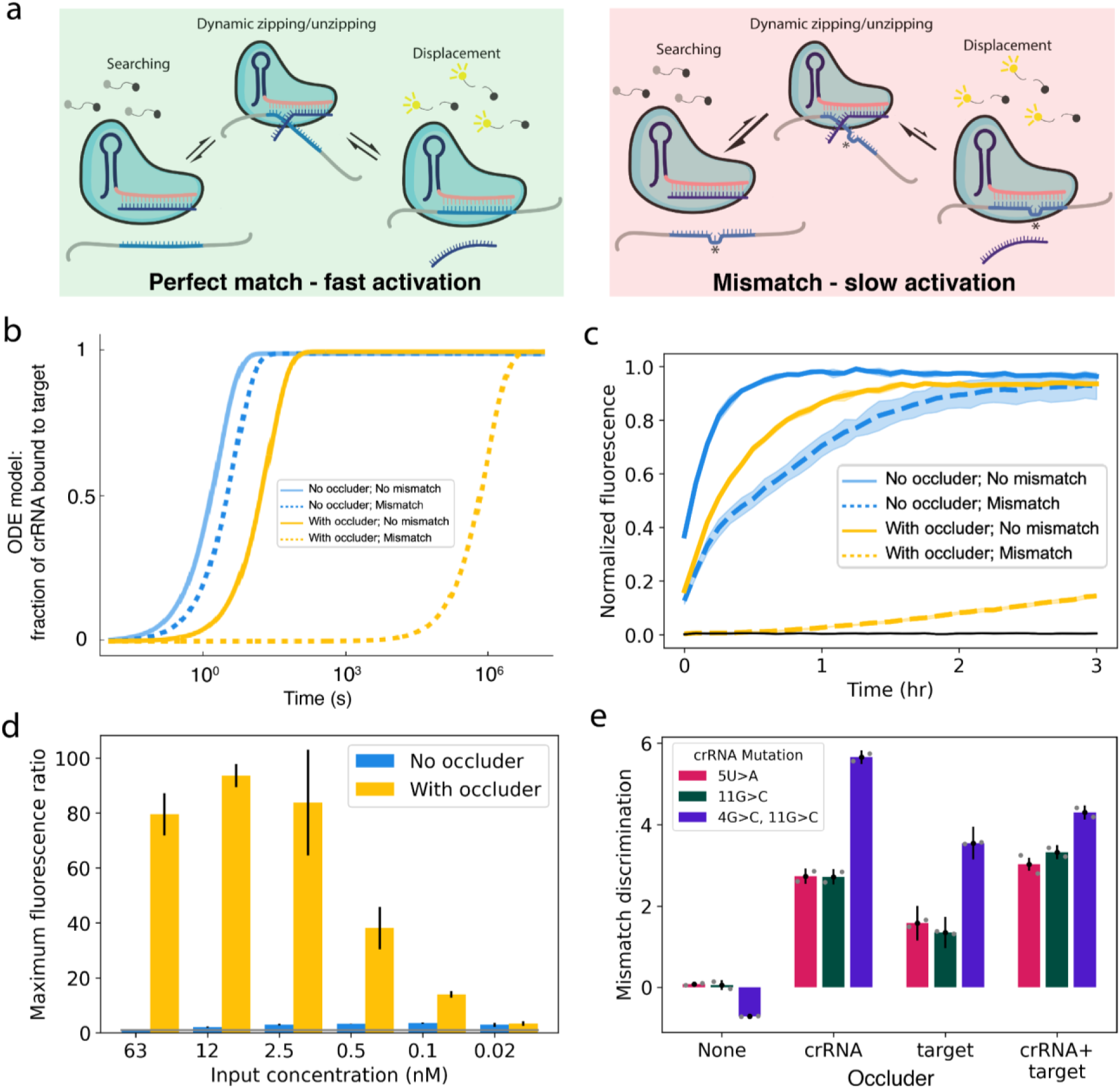
Designed secondary structure enhances Cas13 mismatch detection. **A:** Schematic showing strand displacement by Cas13 with a perfectly-matched target sequence versus one containing a mismatch. **B:** ODE-based model predictions of crRNA/target hybridization kinetics with and without occlusion and mismatches. **C:** Kinetic curves showing detection of a target sequence with and without a single A>U mismatch at spacer position 5, in the presence and absence of occlusion. Shaded region shows range of fluorescence measurements for each condition across replicates. **D:** Maximum fluorescence ratios with and without occlusion at a variety of target input concentrations (see Methods). **E:** Mismatch discrimination, defined as log_2_(perfect match activity / mismatch activity), for two single mutations and one double mutation using occluders blocking either the target, the crRNA, or both. Error bars in D and E show standard deviations propagated through the relevant formulae (see Methods).

We tested Cas13’s ability to differentiate between a perfectly complementary target and one containing a single A>U mutation at position 5 of the protospacer with and without secondary structure occlusion. We tested both occluding the target (as we did previously) and occluding the crRNA, reasoning that strand displacement would in either case result in improved mismatch discrimination. The presence of a crRNA occluder resulted in a ∼50x enhancement of specificity compared to the no-occluder condition, measured as the maximum ratio of WT/mismatch fluorescence (Fig. 3C). This effect was robust to large variations in concentration, and was maximized at higher ∼1-100 nM target input concentrations (Fig. 3D, Extended Data Fig. 7B).

After testing target-occluding and crRNA-occluding oligos, and combinations of both (Fig. 3E, Extended Data Fig. 5), we decided to focus on crRNA occluders. These provide the added benefit of improving mismatch detection regardless of the identity of the mismatched target, and they do not require any sample manipulation prior to detection.

We proceeded to explore the generality of Cas13 specificity enhancement by occluders. Using 3 different targets, 4 positions on each crRNA, and 2 mutations for each position, we tested how well a mismatch could be detected by Cas13 with and without a crRNA-occluder. We measured Cas13 activity on the perfectly matched and mismatched targets, finding that although only 74/96 mismatches (77%) led to any activity reduction in the absence of a crRNA-occluder, all 96 (100%) led to a reduction with the occluder (Fig. 4A-B, Extended Data Fig. 6). Error was thus reduced from 23% to <1%. We found that without occlusion, the ability of Cas13 to distinguish between a perfectly-matched target and a mismatched target is not guaranteed for any crRNA, for any of the mismatch positions we tested, nor for any specific type of mutation (with the exception of G>U, which has the fewest data points). However, using occluded crRNAs, Cas13 can distinguish perfectly-matched from mismatched targets in a position- and mutation-independent manner.

**Fig. 4:**
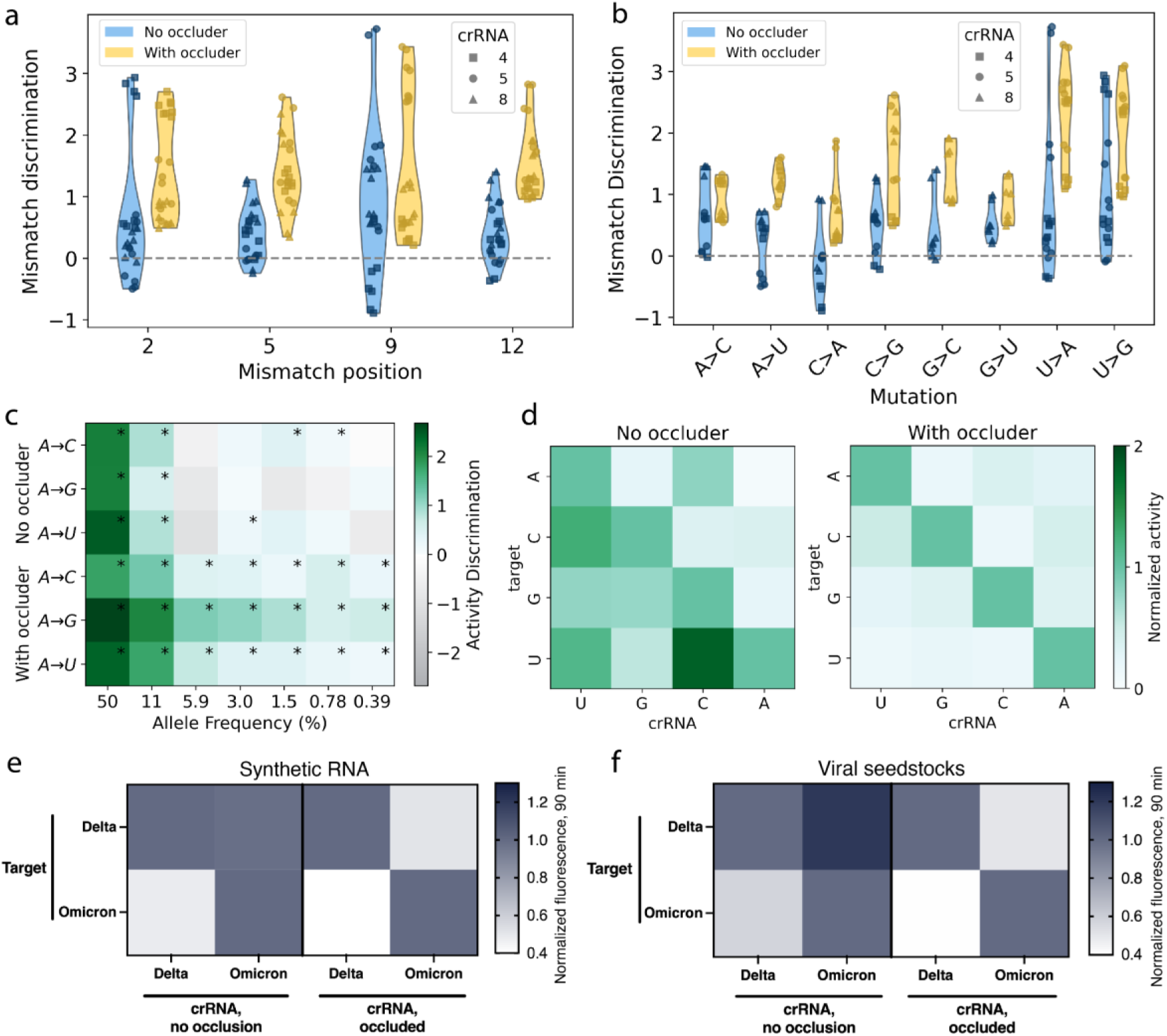
crRNA occluders enable consistent and sensitive mismatch discrimination. **A:** Violin plots showing the ability of Cas13 to distinguish between wildtype targets and targets containing mutations at four different positions in the protospacer both with and without occlusion; position is relative to the 5’ end of the protospacer. Each data point is the discrimination ratio of perfect-match to mismatched sequence (defined as in Fig. 3E; see Methods). **B:** Data from A, but organized by mutation type. **C:** Heatmap showing the ability of Cas13 to detect spiked-in target in a background of mismatched sequence at decreasing allele frequencies, both with and without occlusion. Asterisks indicate statistically significant detection over the no-spike in control. Significance determined by one-tailed t-test p<0.05. Activity discrimination is defined analogously to mismatch discrimination (see Methods). **D:** Specificity matrix showing Cas13 activity normalized for each target, with and without occlusion, for all possible crRNA and target nucleotides at position 5. **E:** Detection of synthetic Delta and Omicron SARS-CoV-2 Spike gene RNA with crRNAs specific to Delta or Omicron using total fluorescence after 90 minutes. Fluorescence of each target was normalized independently after subtracting minimum NTC at 90 minutes. **F:** Same as in E, but using amplified viral seedstocks.

We performed a dilution series of perfectly matched target RNA and mixed it with background RNA containing a single mismatch. Without occlusion, the perfectly matched target was detected at allele frequencies of 11% (a 1:8 ratio) but not 6% (1:16); with occlusion, it was detected at frequencies as low as 0.4% (1:256) for all tested targets, an order-of-magnitude sensitivity enhancement (Fig. 4C, Extended Data Fig. 7A).

To explore nucleotide-specific effects, we mutated the crRNA and target sequences to all possible nucleotides at position 5 of the spacer. When testing all pairwise crRNA/target combinations, we observed extensive cross-reactivity between crRNAs and targets in the absence of occlusion. With occlusion, we observed specific detection of each target only by its perfect-match crRNA (Fig. 4D, Extended Data Fig. 8). Occluded crRNAs can thus discriminate all four possible alleles at a given position, demonstrating the exquisite specificity of the approach.

To test the efficacy of our occlusion strategy in a real-world detection scenario, we designed crRNAs targeting a single locus in the SARS-CoV-2 spike gene containing lineage-specific mutations in adjacent codons. In this way, it is theoretically possible to distinguish Delta and Omicron strains with only a single crRNA designed to identify each strain. Targeting polymorphic sequences with conventional Cas13 detection schemes is challenging, as crRNAs targeting a given mutation are likely to be affected by nearby mutations, making it nearly impossible to distinguish variants from one another. We found that with crRNA occlusion, but not in its absence, targeting this single locus leads to specific detection of the target strain in both synthetic RNA (Fig. 4E, Extended Data Fig. 9) and amplified viral seedstocks (Fig. 4F) using a simple fluorescence readout.

## Discussion

In this study, we quantified the reduction in Cas13 activity due to target RNA occlusion, showed that our results are quantitatively consistent with a strand-displacement-based model of Cas13 activation, and used this model to improve Cas13’s mismatch specificity by an order of magnitude.

Importantly, our method is simple to implement experimentally as it merely requires annealing a DNA oligo to the crRNA prior to mixing with the target RNA. Unlike in other studies leveraging RNA secondary structure to improve hybridization specificity, no toehold is required, enabling trivial construction of the DNA oligo: all crRNA occluders used occluded the entire crRNA spacer. The main downside of using crRNA-occluder duplexes to improve mismatch detection is a reduction in Cas13 activity. To counteract this, steps can be taken to increase overall activity such as increasing concentrations of Cas13, reporter, or target RNA.

While our experimental study focused on one ortholog of Cas13, LwaCas13a, our results are of interest to many Cas13-based methods. Future work will explore to what extent the mechanism of activation described here is generic or specific to this ortholog. Furthermore, significant questions remain regarding the mechanism by which secondary structure 3’ to the protospacer leads to an allosteric reduction in Cas13 activity. For instance, extended complementarity between the target and the structural portion of the crRNA (known as the the tag-antitag effect) inhibits Cas13 activity by preventing the proper formation of the active site^36,37^. Future structural and biochemical characterization will reveal whether the inhibitory region we have discovered affects Cas13 by a similar or a different mechanism.

Due to its negligible cost, ease of implementation, orthogonality with existing approaches, and marked improvement in detection specificity, we anticipate the adoption of our crRNA occlusion approach into a wide range of Cas13-based techniques. Furthermore, our proposed strand displacement model addresses a long-standing paradox, namely how a purportedly ssRNA-specific enzyme is able to robustly target RNAs in cellular environments where RNA structure is ubiquitous.

## Methods

### General reagents

Oligonucleotides were ordered from Integrated DNA Technologies (IDT). Unless otherwise noted, chemical reagents were ordered from Sigma. Oligonucleotide sequences are listed in Supplementary File 1.

### crRNA design

Most crRNA spacers were designed to be perfectly complementary to their 28 nucleotide (nt) protospacer region. For SARS-CoV-2 targeting sequences, a single synthetic mismatch was inserted at position 5 to improve baseline specificity. Spacers were appended to the 3’ end of the consensus LwaCas13a direct repeat sequence (AGACUACCCCAAAAACGAAGGGGACUAAAAC) and ordered from IDT as Alt-R guide RNA.

### Target design

For our tiling experiment, we designed an RNA molecule of length 961 nucleotides with minimal internal secondary structure. After an initial G nucleotide, the molecule is comprised of 10 target blocks, each defined by a 34-nt buffer region, a 28-nt protospacer, and a second 34-nt buffer region. We sought to have as many as possible of the 28-nt protospacers resemble natural sequences.

To this end, we started with a set of 18,508 28-nt-long protospacer sequences compiled from the ADAPT dataset, which has a sequence composition representative of viral diversity^12^. 3,391 sequences with poly-A, poly-C, or poly-U stretches ≥5 nts or poly-G stretches ≥4 nts were removed. Of the remaining sequences, we removed 6,459 which had low average measured activity, defined as *〈*out_log_k〉≤ -2 (on a logarithmic scale from -4 to 0, where 0 is high activity) using the activity definitions and measurements from Ref^12^. We used LandscapeFold^38^ with parameter 𝑚 = 2 (𝑚 represents the minimum allowed stem length), disallowing pseudoknots, to predict the structure landscapes of the remaining sequences. LandscapeFold predicted that 1,287 of these remaining sequences had extremely low intramolecular structure, defined as all nucleotides having a ≥40% probability of being unpaired in equilibrium.

We then aimed to find a set of these sequences that were all dissimilar from one another. First, given a sequence 𝑠, we found all those sequences with a Hamming distance ≤15 from 𝑠. (A pair of sequences with a Hamming distance of ℎ share all but ℎ nucleotides). Of these sequences, we chose the one with the least secondary structure to keep and removed the others, with total amount of secondary structure quantified as 𝛴_𝑛_𝑝_𝑛_ where the sum is over nucleotides and 𝑝_𝑛_ is the probability of the nucleotide being paired in equilibrium. We repeated this step for each sequence 𝑠 we had not already removed. Next, we used a Smith-Waterman alignment^39^ to check for sequence similarity in non-identical nucleotide positions, repeating the same procedure as above but, instead of Hamming distance, using the criteria of an alignment score ≥9 to define sequence similarity, where the alignment score parameters were (+1, -2, -2) for (match, mismatch, gap). This procedure resulted in a set of 20 sequences all distant from one another in sequence space.

Finally, although we had ensured each of these sequences had low secondary structure, we wanted to minimize binding between these sequences. For each pair of sequences, we used LandscapeFold with parameter 𝑚 = 3 to predict the structure of the two strands, allowing for both intra- and inter-molecular interactions. We defined two sequences to be incompatible if the resulting prediction had any nucleotide on either sequence with a ≤40% probability of being unpaired in equilibrium. We exhaustively enumerated the possible ordered sets of mutually compatible sequences, finding 60 ordered sets of 5 mutually compatible sequences, and no set of 6 mutually compatible sequences. Of these 60 sets, we chose the one with the least structure. Under the assumption that entropic loop closure costs will create a barrier to non-neighbor sequence pairing (i.e. that each sequence is less likely to pair to a sequence that isn’t its neighbor), we defined structure here as the sum, over the 4 pairs of neighboring sequences, of the maximum probability of a nucleotide being paired in that sequence pair. Thus, we arrived at a set of 5 distinct sequences from ADAPT with minimal intra- and inter-molecular structure. These 5 sequences became the protospacer sequences corresponding to crRNAs 2, 4, 6, 8, 9.

The other 5 protospacer sequences as well as the buffer regions were compiled out of 64 16-nt-long DNA sequences with minimal internal structure from Shortreed *et al.*^27^. Seven of these sequences with poly-A or poly-T stretches ≥5 nts were removed. Concatenating these sequences resulted in a long sequence with minimal structure, which we used to construct the rest of the 961 nt-long RNA target. We used NUPACK 3^40^ to predict the structure of the resulting target, finding various predicted stems. We then made individual point mutations by hand in the buffer regions and non-ADAPT-derived protospacers to minimize the probabilities of the resulting stems (ensuring NUPACK predicted no base pair forming with probability ≥60% in equilibrium), as well as to remove sequence similarity between targets (ensuring there are no more than 5 identical consecutive nucleotides between the protospacer regions, no more than 6 identical consecutive nucleotides between two regions spanning a protospacer and a buffer, and no more than 8 identical consecutive nucleotides in buffer regions).

Finally, we created a “shuffled” version of the target, placing the target blocks (numbered 1 – 10 from 5’ to 3’ in the original target) in the following order: 1, 4, 2, 7, 5, 3, 9, 6, 8, 10. We ensured NUPACK 3 did not predict any base pair forming with probability ≥60% in the resulting sequence.

For our initial experiments (Fig. 1) we filtered the ADAPT dataset sequences to those with high activity (<OUT_LOG_K> -2) and perfect complementarity between target and crRNA in the ADAPT dataset. We then measured LandscapeFold’s prediction of the secondary structure of each candidate protospacer sequence. For each nucleotide, we calculated the total probability that the nucleotide is unpaired in equilibrium. The protospacer chosen had each nucleotide with at least a 92% probability of being unpaired in equilibrium.

### RNA preparation (including structured targets)

RNA targets were ordered from Integrated DNA Technologies as DNA containing a T7 promoter sequence. Targets were then transcribed to RNA using the T7 HiScribe High Yield RNA Synthesis Kit in 55 μL reactions (New England Biolabs) with a 16h incubation step at 37 °C and purified with 1.8X volume AMPure XP beads (Beckman Coulter) with the addition of 1.6X isopropanol, then eluted into 20 μL of nuclease free (NF) water. All RNAs were then quantified using a NanoDrop One (Thermo Fisher Scientific) or Biotek Take3Trio (Agilent) then stored in nuclease free (NF) water at -80 °C for later use.

Occluded targets and crRNAs were prepared by mixing DNA/RNA oligo occluders with target RNA or crRNA in 60mM KCl (Invitrogen) in NF water at a ratio of 2:1 (BioMark assays) or 10:1 (plate reader assays) and put through an annealing cycle consisting of a high-temperature melting step at 85 °C for three minutes followed by gradual cooling to 10 °C at 0.1 °C/sec followed by cooling to 4 °C. For massively multiplexed assays, occluders were first pooled by length and start position within the target block (see Extended Data Fig. 1D) such that each resulting oligo pool contained all 8 n-mers binding to a given position within each of the experimental target blocks. Targets and crRNAs were then used for detection assays immediately as described below.

Targets were input into detection reactions at various concentrations. For experiments in Fig. 1, targets were input at 7.5 x 10^8^ copies/μL (cp/μL). For experiments in Fig. 2a-e, targets were input at 8 x 10^8^ cp/μL. For Fig. 2f, targets were input at 5 x 10^9^ cp/μL. For experiments in Fig. 3 and 4a-b, input concentrations of 7.5 x 10^9^ cp/μL were used unless otherwise noted in figure caption. For Fig. 4c, targets were spiked in at the indicated allele frequency into a background of 5 x 10^10^ cp/μL (for occluded conditions) or 5 x 10^8^ cp/μL (for non-occluded conditions). For Fig. 4d, occluded conditions used an input concentration of 5 x 10^10^ cp/μL while non-occluded conditions used a concentration of 5 x 10^8^ cp/μL.

### Viral seedstock amplification

Extracted viral genomic RNA samples were acquired from BEI Resources (hCoV-19/USA/MD-HP05285/2021 (B.1.617.2) Delta, hCoV-19/USA/GA-EHC-2811C/2021 (B.1.1.529) Omicron).

Amplification reactions using 2uL of viral RNA as the input (50uL total reaction volume) were performed using the Qiagen One-Step RT-PCR kit according to the manufacturer’s specifications. The forward primer contained a T7 promoter sequence; 4uL of the RT-PCR products were used as direct input for T7 transcription (see “RNA preparation” section above).

### Cas13 detection assays

Standard bulk detection assays were performed by mixing target RNA at a ratio of 10% v/v with 90% Cas13 detection mix. The detection mix consisted of 1X RNA Detection Buffer (20 mM HEPES pH 8.0, 54 mM KCl, 3.5% PEG-8000 in NF water), supplemented with 45nM purified LwaCas13a (Genscript, stored in 100 mM Tris HCl pH 7.5 and 1 mM DTT), 1 U/μL murine RNAse Inhibitor (New England Biolabs), 62.5 nM fluorescent reporter (/5FAM/rUrUrUrUrUrU/IABkFQ/; IDT), 22.5 nM processed crRNA (IDT), and 14 mM MgOAc. In experiments using crRNA occlusion, crRNAs were pre-annealed to DNA occluders as described above, and used at a final concentration of 22.5 nM. 15 μL reactions were loaded in technical triplicate onto a Greiner 384 well clear-bottom microplate (item no. 788096) and measured on an Agilent BioTek Cytation 5 microplate reader for 3 hours with excitation at 485 nm and detection at 528 nm every five minutes.

For tiling assays, Standard Biotools genotyping IFC (192.24 format) was used in a BioMark HD for multiplexed detection. Assay mix (10% of final reaction volume) contained 1X Assay Detection Mix (Standard Biotools) supplemented with 100 nM crRNA, 100 nM LwaCas13a (Genscript, stored in 100 mM Tris HCl pH 7.5 and 1 mM DTT). Sample mix (90% of final reaction volume) contained 1X Sample Buffer (44 mM Tris-HCl pH 7.5, 5.6 mM NaCl, 10 mM MgCl, 1.1 mM DTT, 1.1% w/v PEG-8000), supplemented with murine RNAse Inhibitor (1 U/μL, NEB), fluorescent reporter (500 nM, IDT), 1x ROX reference dye (used for normalization of random fluctuations in fluorescence between chambers) (Standard Biotools), 1X GE Buffer (Standard Biotools), 20mM KCl, and occluded RNA target (9 x 10^8^ cp/μL).

Sample volumes of 3.5μL and assay volumes of 3.5μL, in addition to appropriate volumes of Control Line Fluid, Actuation Fluid, and Pressure Fluid (Standard Biotools), were loaded onto 192.24 genotyping IFC chip (Standard Biotools). Chips were then placed into the Fluidigm Controller and loaded and mixed using the Load Mix 192.24 GE script (Standard Biotools).

After mixing, reactions were run on BioMark HD at 37°C for 8 hours with measurements taken in the fluorescein amidite (FAM) and the carboxyrhodamine (ROX) channels every five minutes.

Normalized and background-subtracted fluorescence for a given time point was calculated as (FAM - FAM_background) / (ROX - ROX_background) where FAM_background and ROX_background are the FAM and ROX background measurements.

### Activity fits

Fluorescence curves were converted to activity scores by fitting the curves to effectively first-order reactions. With a certain amount of active Cas13, the concentration of uncleaved reporter is expected to decrease exponentially according to the reaction

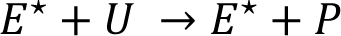

where 𝐸^⋆^ is the concentration of active Cas13, 𝑈 the concentration of uncleaved reporter, and 𝑃 the concentration of cleaved reporter RNA. Labeling the (second-order) rate constant of this reaction as 𝑟, the concentration of 𝑃 changes over time according to

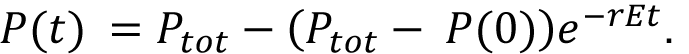

Assuming that 𝐸^⋆^ is constant over time, we define an activity score 𝜈 = 𝑟𝐸^⋆^, which is an effective first-order rate constant (with units of inverse time). Assuming that 𝑟 is constant across our assays, the activity score 𝜈 is thus a proxy for the amount of active Cas13. Given measured P(0), we find best-fit values of P_tot_ and 𝜈 to fit the kinetic curves. To account for curves very far from saturation (e.g. NTC data) we set a minimum value of P_tot_ based on data from saturating and near-saturating curves. For tiling data, we fit the first 50 timepoints (∼4 hours) to discount occasional apparent noise appearing at very late times.

Some assays including crRNA occluders displayed fluorescence curves that did not fit well to this effective first-order reaction (Extended Data Figs. 5, 6, 7, 8) indicating a need to relax the assumption that 𝐴 is constant over time. For data shown in Extended Data Figs. 7A and 8, we neglected the first several timepoints measured (15 and 10 timepoints, respectively, corresponding to 75 and 50 min) since we found that doing so increased the goodness of fit. For other assays using crRNA occluders—and those assays being directly compared to them (i.e. those data shown in Extended Data Figs. 5, 6, 7B)---we fit the data to a series of two effective first-order reactions:

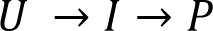

Labeling the first-order rate constant of each reaction 𝑘_1_ and 𝑘_2_, this model yields

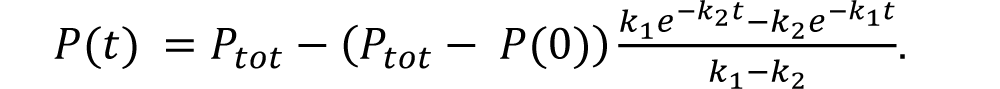

We define activity in this case as 𝜈 = (1 / k_1_ + 1 / k_2_)^-1^, verifying that if the equation is first order (i.e. k_1_ ≫ k_2_), our previous definition of activity is recovered. We indeed find negligible change in the measured activities for no-occluder control (NOC) fluorescence curves between these two fits.

### Tiling experiment activity correction

Each experimental condition in the tiling experiment was performed with four replicates: two technical replicates for each of the two “shuffles” of the 961-nt-long target sequence. While we found excellent agreement between technical replicates (Extended Data Fig. 3B), there was some variation between the results from each of the two target “shuffles” (Extended Data Fig. 3H). This variation was apparent in and correlated between the positive controls of crRNAs 1 and 10, which were always unoccluded (Extended Data Fig. 3C, H). We hypothesized that this variability results from small variations in target concentration in our different samples.

To correct for such variations, we sought to quantify how much each RNA sample differed from the mean. The RNA samples were divided into 192 sample conditions, each corresponding to a single oligo pool and one target shuffling (see “RNA preparation” section, Extended Data Fig. 1C). Each of these 192 conditions was mixed with 24 assay conditions, corresponding to 8 experimental crRNAs, 2 positive control crRNAs, one non-targeting crRNA, one no-crRNA control (NPC) and two replicates of each.

For the two replicates of crRNAs 1 and 10 (i.e. for each of the 4 positive control assay conditions out of the 24 total assays), we considered the activity fit from the mean fluorescence curve, averaging over the 192 sample conditions (Extended Data Fig. 3B, black dashed lines). Then, for each sample condition, and for each positive control, we calculated the ratio of the control’s activity to its mean activity across all samples, obtaining an estimate of the degree to which that sample’s concentration deviated from the mean. We defined a correction factor as the average of these ratios. We then divided all activities measured for that sample by this correction. This activity correction not only decreased the spread of activities measured by the positive controls (Extended Data Fig. 3D-G) but also decreased the variance between measurements made on the two target “shuffles” (Extended Data Fig. 3H).

### Mismatch discrimination

In Fig. 3D we show the results of one simple metric by which to measure mismatch discrimination: the maximum of the ratio F_PM_ / F_MM_, where F_PM_ is the average fluorescence measurement across the perfectly matched conditions, and F_MM_ across the mismatched conditions. To account for the arbitrary offset of fluorescence, the minimum fluorescence measured across the NTC experiments was subtracted from both F_PM_ and F_MM_ before taking the ratio. Since F_PM_ and F_MM_ are each measured as the average across three replicates, each measurement of F_PM_ and F_MM_ has an inherent error, σ_PM_ and σ_MM_ (respectively), which we quantify as the standard deviation across the three replicates at each timepoint. The error of the ratio is then propagated as

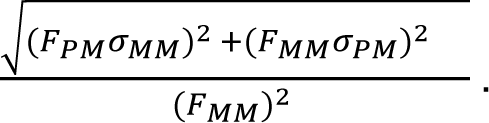

Elsewhere (Fig. 3E, Figs. 4A, B) we measure mismatch discrimination by a metric that relies on activity fits: log_2_(𝜈_PM_ / 𝜈_MM_). Thus, a mismatch discrimination of 1 indicates that the measured activity of the perfectly matched conditions is twice that of the mismatched conditions, and a discrimination of 3 indicates the perfectly matched conditions had eight-fold higher activity than the mismatched conditions. We used a similar measure for discrimination at low allele frequencies (Fig. 4C), defining activity discrimination as log_2_(𝜈_f_ / 𝜈_0_) where 𝜈_f_ is the activity measured at allele frequency *f*, and 𝜈_0_ the activity measured in the background alone.

### Equilibrium model

In an equilibrium model of crRNA-target hybridization, the target has a free energy ΔG_u_ that depends on occluder conditions, and the crRNA-target complex has free energy independent of occluder. A kinetic model describing crRNA-target hybridization would be

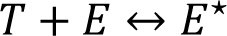

with the ratio of forward to reverse rate constants determined by the free energy difference between the target-unbound and target-bound states. Assuming that the target is in excess, we define an unknown parameter 𝛼_1_ such that at steady state,

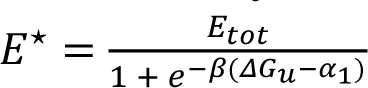

where β = 1 / k_B_T, with k_B_ denoting Boltzmann’s constant and T temperature (measured in Kelvin), and E_tot_ = E + E* is the total concentration of Cas13 enzyme in the system (measured in Molar).

Our measured activity, 𝜈, is proportional to E* as defined previously: 𝜈 = rE*. Considering 𝜈 on a logarithmic scale (as it is plotted in Fig. 1C) yields

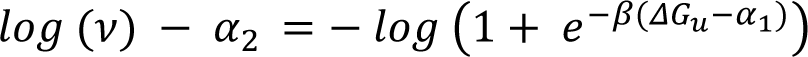

with 𝛼_2_ =𝑙𝑜𝑔 (𝑟𝐸_𝑡𝑜𝑡_).

The two parameters ɑ_1_ and ɑ_2_ therefore only shift the curve relating 𝑙𝑜𝑔 (𝜈) to ΔG_u_, but cannot alter the shape of the curve itself.

That the relationship between 𝑙𝑜𝑔 (𝜈) and ΔG_u_ is approximately linear can be seen by recognizing that if

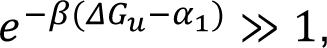

as is typical in our system, we have

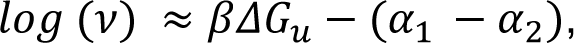

indicating that there is in essence only one free parameter in the system that shifts the line but cannot change its slope. The slope is set by the temperature of the system.

### ODE model

We compared the binding rate of an invading strand to a target with and without an occluder in a model based on Srinivas & Oulridge *et al.* (2013) and Irmisch *et al.* (2020)^29,35^. The model consists of a set of ordinary differential equations (ODEs) representing the flux into and out of states, where each state is defined by the set of base pairs formed. Transitions between states occur at a rate ke^-ΔG^ where ΔG is the free energy barrier to the transition (in units of k_B_T where k_B_ is Boltzmann’s constant and T is temperature in units of Kelvin) and 𝑘 is an overall rate constant. An initial state consists of a target strand (bound to an occluding strand in the case where an occluding strand is considered), with an invading strand unbound. Initial binding of the invader strand to the toehold has a free energy barrier of ΔG_a_. The reverse step has a barrier of hΔG_R_ where ℎ is the toehold length and ΔG_a_ is the (absolute value of the) typical free energy of an RNA-RNA base pair.

In the no-occluder case, subsequent forward steps (in which an additional base pair between target and invader forms) have a free energy barrier of 0, while reverse steps have a free energy barrier of ΔG_a_.

In the occluder case, the first step of the strand displacement reaction has barrier ΔG_P_ + ΔG_S_ − (ΔG_R_ − ΔG_D_), where we have subtracted (ΔG_R_ - ΔG_D_) from the models on which we base our work to account for the fact that in our system, the invading strand is RNA while the occluding strand is DNA; ΔG_D_ is the (absolute value of the) typical free energy of an RNA-DNA base pair. Subsequent forward steps in the strand displacement reaction have a barrier of ΔG_S_ − (ΔG_R_ −ΔG_D_), while backward steps all have a barrier of ΔG_S_. The barrier from the final state, in which the occluder has fully dissociated, back to the penultimate state, has a barrier of ΔG_D_.

Parameters were set following Irmisch *et al.* to: ΔG_a_ = 18.6; ΔG_R_ = 2.52; ΔG_s_ = 7.4; ΔG_p_ = 3.5; ΔG_m_ = 9.5 (all in units of k_B_T); and an overall rate constant of 𝑘 = 6 x 10^7^/s. We set ΔG_D_ = 1.2 to be roughly half of ΔG_R_, and ΔG_D_ = 25 to be large enough to prevent reassociation on the timescales considered. In Fig. 3C we plot the results of *h* = 3, *b* = 27, with a mutation at the first position after the toehold.

### Strand displacement model

In the strand displacement model, non-specific binding of Cas13 to the RNA (independent of RNA sequence) leads to Cas13 activation, and a corresponding decrease in Cas13 dissociation rate, when the crRNA fully binds to the protospacer complement. We denote the typical dwell time of Cas13 on the RNA in the absence of this activation by t_dwell_, the main parameter in the model. Secondary structure affects activity by modulating the probability that a strand displacement reaction completes within this time t_dwell_. We assume that activity is directly proportional to this probability. To estimate this probability, for each occluder, we simulated 10^5^ unbiased random walks of length equal to the number of occluded protospacer nucleotides with a reflecting boundary at 0, measuring the number of steps taken to complete the random walk, and therefore the probability of completing the random walk within a desired number of steps.

The number of trials chosen leads to errors in our estimate of this probability < 3% in all cases (determined by the maximum ratio of standard deviation to the mean of our estimate across 10 replicates of 10^5^ random walks).

To estimate a typical value of t_dwell_, we turn to classic studies of the *E. coli lac* repressor, a DNA-binding enzyme which searches for its binding site on the DNA by iteratively binding non-specifically to the DNA, performing a local search, and dissociating^25,26^. The dissociation rate of the *lac* repressor when non-specifically binding DNA (i.e. the inverse of its dwell time) has been estimated^25^ to be 5 x 10^4^/s. This dwell time corresponds to the time it would take for ∼100 steps of a strand displacement reaction, where the rate of individual steps has been estimated^35^ to be 6 x 10^7^ x e^-2.5^/s ≈ 5 x 10^6^/s. This dwell time varies with ionic concentration (among other factors), with dwell time decreasing anywhere from 2-10-fold upon doubling KCl concentration^25^. Given the different conditions used in the plate-reader assays and in the tiling experiments, including different ionic conditions—with the former having ∼2.5-fold higher KCl concentrations than the latter—we fit t_dwell_ separately for the two experimental methods. We used a dwell time of 100 steps of a strand displacement reaction for the plate-reader assays, and a dwell time of 300 steps for the tiling experiments.

To account for the asymmetry seen in our data (Fig. 2D, Extended Data Fig. 4A-D) we allowed for dwell time to change depending on whether the 5’ end of the protospacer was occluded or unoccluded. For the plate reader assays, our final model has a dwell time of 100 steps for those cases where the 5’ end of the protospacer was occluded, and a dwell time of 200 steps for those cases where it wasn’t. For the tiling assays, the dwell times used are 300 and 600 steps, respectively.

## Acknowledgments

We thank Brian Kang, Britt Adamson, Rees Garmann, and Herman Dhaliwal, as well as the members of the Myhrvold and te Velthuis labs, for useful discussions.

## Author contributions

All authors (O.K., B.B.L., O.R.S.D., A.J.W.t.V., C.M) designed research. B.B.L. and O.R.S.D. performed experiments. O.K., B.B.L., O.R.S.D., and C.M. performed data analysis. O.K. and C.M. performed modeling. All authors (O.K., B.B.L., O.R.S.D., A.J.W.t.V., C.M) wrote the article.

## Extended data figures

**Extended Data Fig. 1:**
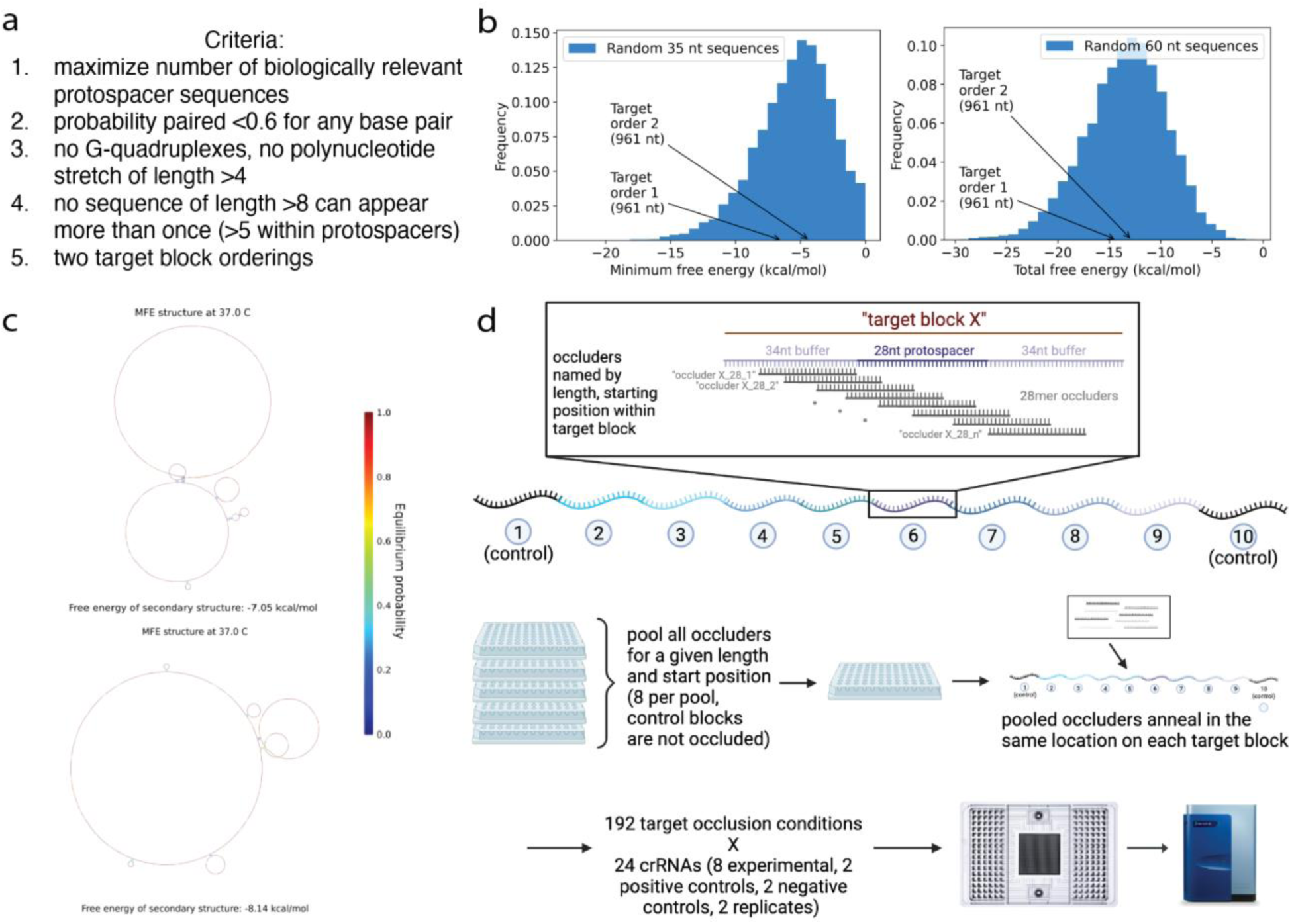
Multiplexing RNA secondary structure for Cas13-based assays. **A.** Criteria used in the design of target RNAs for multiplexed detection assays. **B.** Histogram showing the minimum and total structure free energies of randomly generated RNA sequences, compared to the free energy of the two target RNAs used in the tiling experiment. **C.** NUPACK 3 predicted minimum free energy structures for the two experimental targets^40^. **D.** Overview of the multiplexed tiling experiment. Made using BioRender.

**Extended Data Fig. 2:**
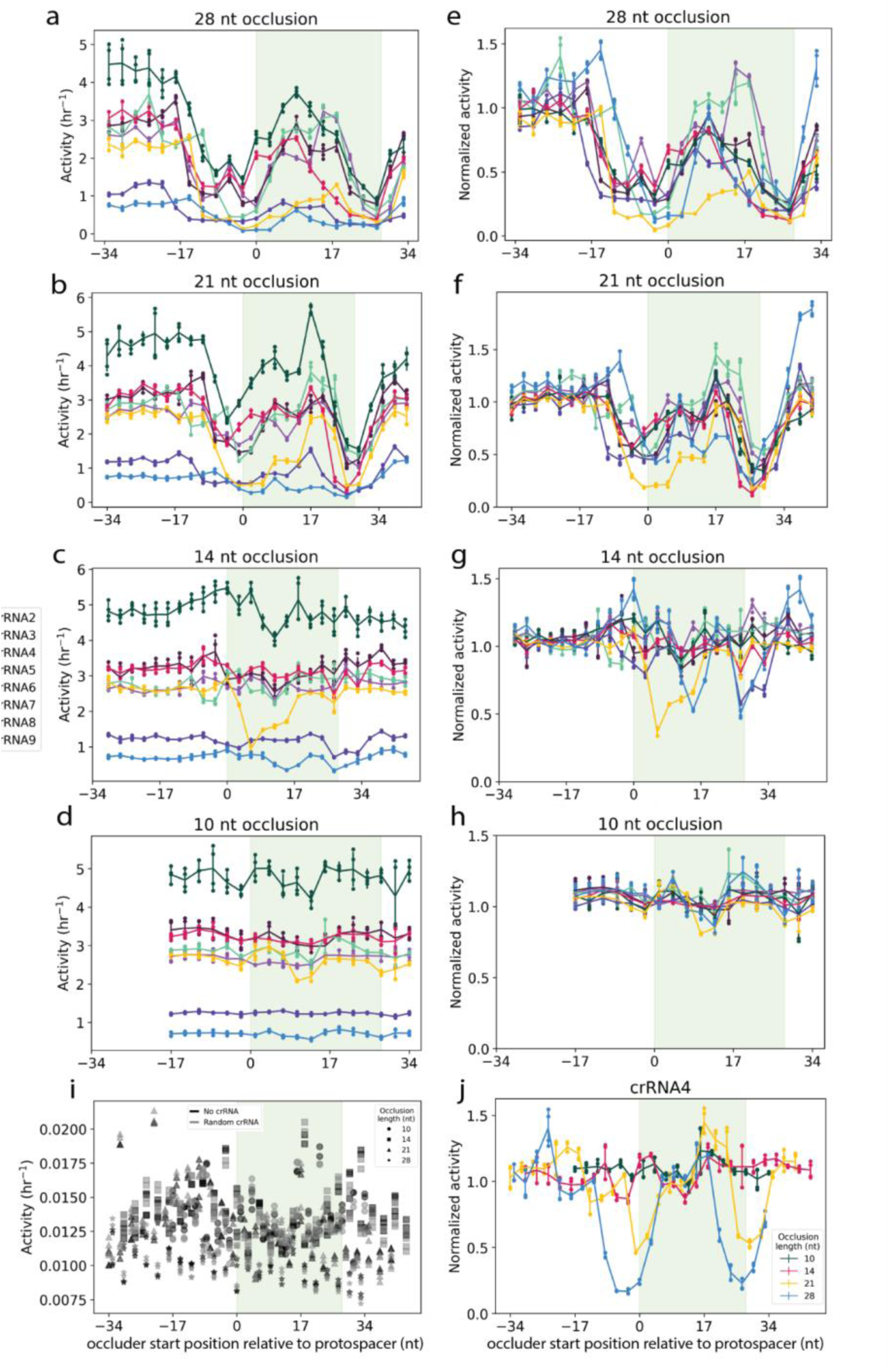
Activity profiles of Cas13 targeting occluded RNAs. **A-D.** Cas13 activity as a function of occluder start position in each 96 nt target block. **A:** 28-mers. **B:** 21-mers. **C:** 14-mers. **D:** 10-mers. **E-H.** Same as A-D, with activities normalized to the non-occluded condition for each target block **I.** Activities of all negative control conditions. **J.** Normalized activities as a function of occluder start position for a single target with all four occlusion lengths.

**Extended Data Fig. 3:**
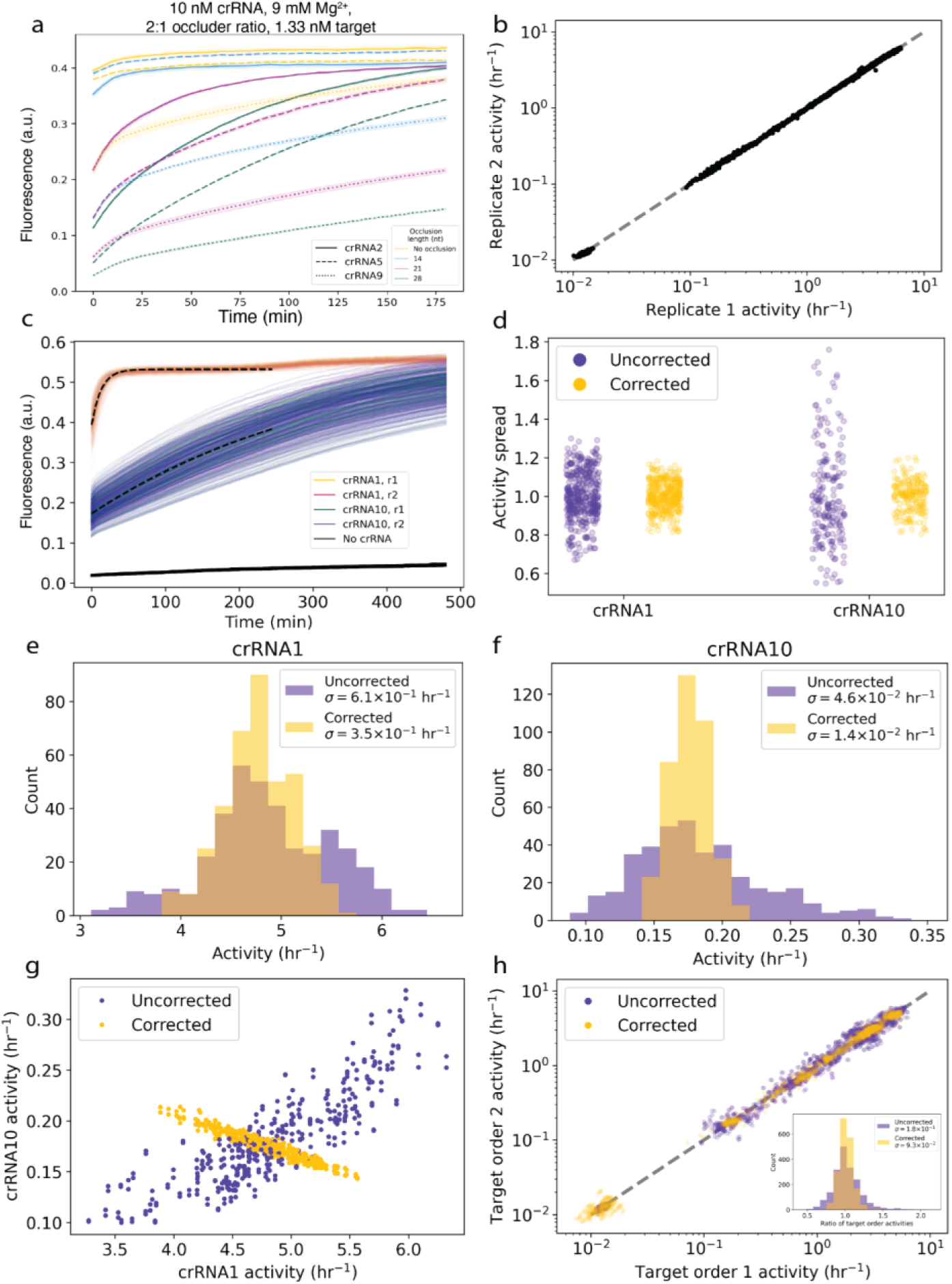
Controls and data normalization for multiplexed tiling assay. **A.** Fluorescence curves for the set of experimental conditions chosen for the multiplexed tiling assay. **B.** Scatter plot showing activity correlation between technical replicates in the tiling experiment. **C.** Raw activity curves from the control (unoccluded) targets across all conditions in tiling assay. **D.** Swarm plot showing control target activities before and after correction. **E.** Cumulative histogram showing activity distribution for control target 1 before and after correction. **F.** Same as in E, but for control target 10. **G.** Dot plot showing activity correlations between the two control targets for all conditions before and after correction. **H.** Scatter plot showing activity correlation between the two target shufflings for all tested conditions, before and after correction. Inset: histogram of ratio of activities of points of the two target shufflings, before and after correction, excluding negative controls. In E, F, H (inset), 𝜎 represents standard deviation.

**Extended Data Fig. 4:**
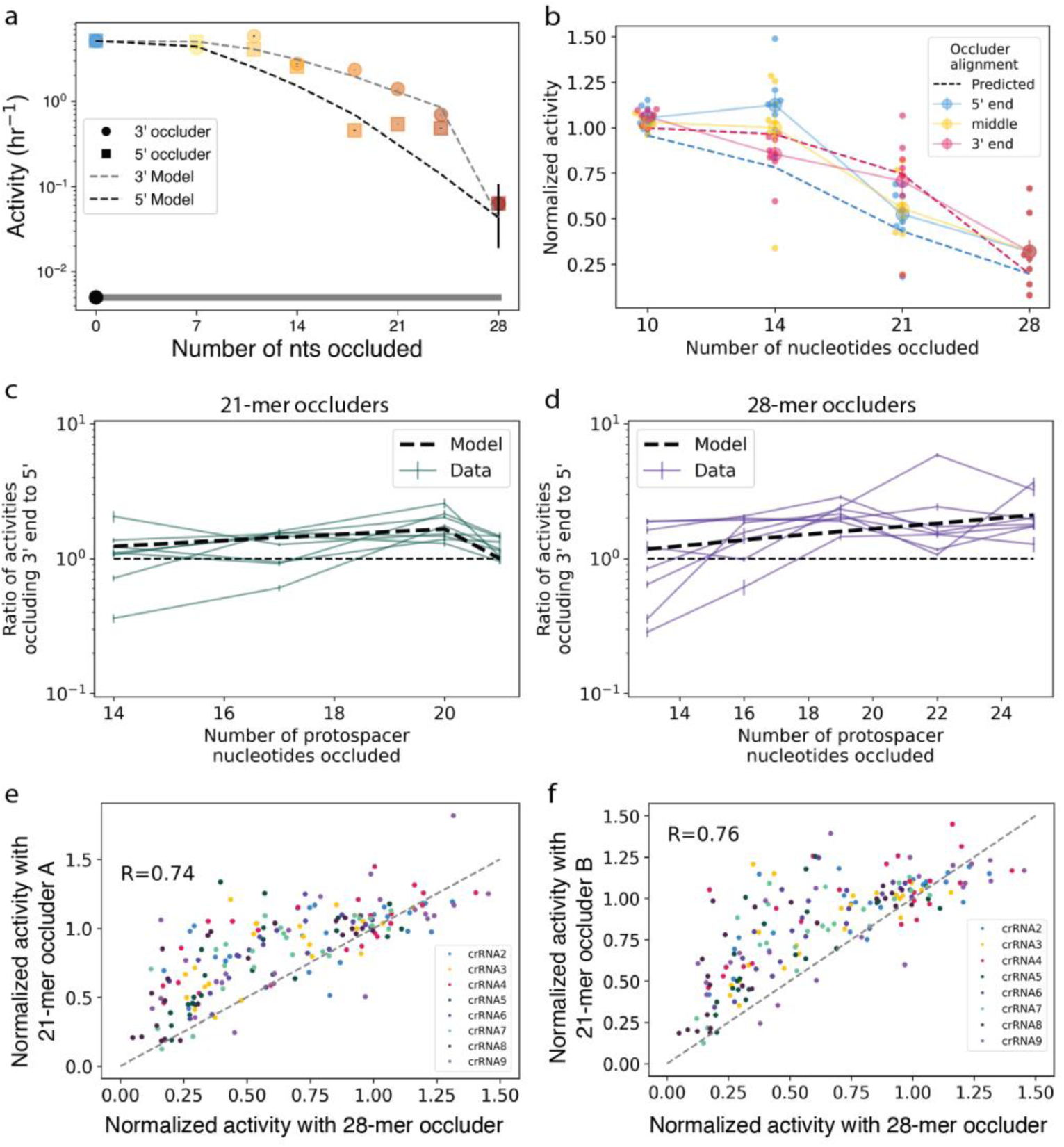
Strand displacement-based t_dwell_ model accounts for asymmetries in occlusion pattern. **A.** Activity as a function of DNA occluder length for occluders of different lengths extending inwards from 3’ and 5’ ends of the protospacer. Strand displacement model predictions shown as dashed lines. **B.** Data from tiling experiment plotted as in A and also including oligos occluding the middle of the protospacer (yellow). Small dots show individual data points from each crRNA; large dots show the mean; error bars show standard deviation. Model predictions shown as dashed lines. **C.** Ratio of activities resulting from 21-mer occluders targeting 3’ end of protospacer to those resulting from occluders targeting 5’ end, plotted as a function of occluder length. Each curve shows a different crRNA. Model prediction shown as dashed line. Cumulative histograms of these data are shown in Fig. 2D. **D.** As in C, for 28-mer occluders. **E-F.** Correlation between activities resulting from 21-mer occluders and 28-mers occluding the same nucleotides. In E, 28-mers starting at positions 1, 4, 7, etc are matched with 21-mers starting at positions 3, 6, 9, etc; in F they are matched with 21-mers starting at positions 6, 9, 12, etc.

**Extended Data Fig. 5:**
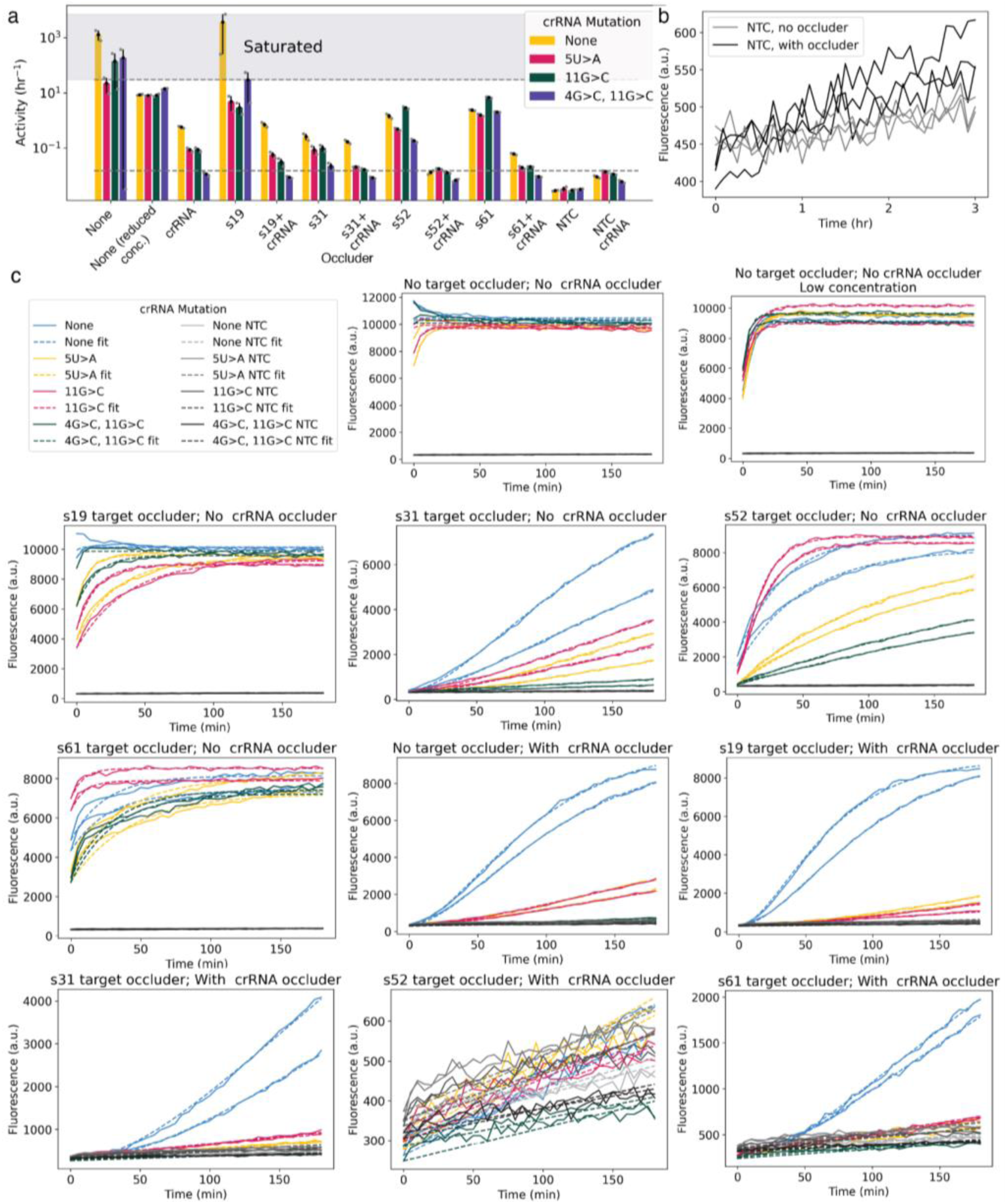
Occlusion of target RNA and crRNA increase Cas13’s sensitivity to mismatches. Note on occluder nomenclature: occluders are named according to their start position from the 5’ end of the target block; all occluders here are 28nt in length. Occluder s19 binds to nucleotides 1-13 of the protospacer and extends into the 5’ flanking region, s31 binds to nucleotides 1-25 and extends slightly into the 5’ flanking region, s52 binds to nucleotides 18-28 and extends into the 3’ flanking region, and s61 binds to nucleotide 28 of the protospacer and extends into the 3’ flanking region. **A.** Bar charts corresponding to Fig. 3E, representing Cas13 activity for perfectly-matched and mismatched targets when occluded by several different target-blocking occluders, as well as crRNA occluders and combinations thereof. Upper dotted line indicates activity saturation point; curves above saturation are not fit well by exponentials and therefore bars in this region are not accurate measurements of activity. **B.** NTC (no-target control) raw curves for conditions with and without a crRNA occluder. **C.** Raw fluorescence curves and fits for the data shown in A.

**Extended Data Fig. 6:**
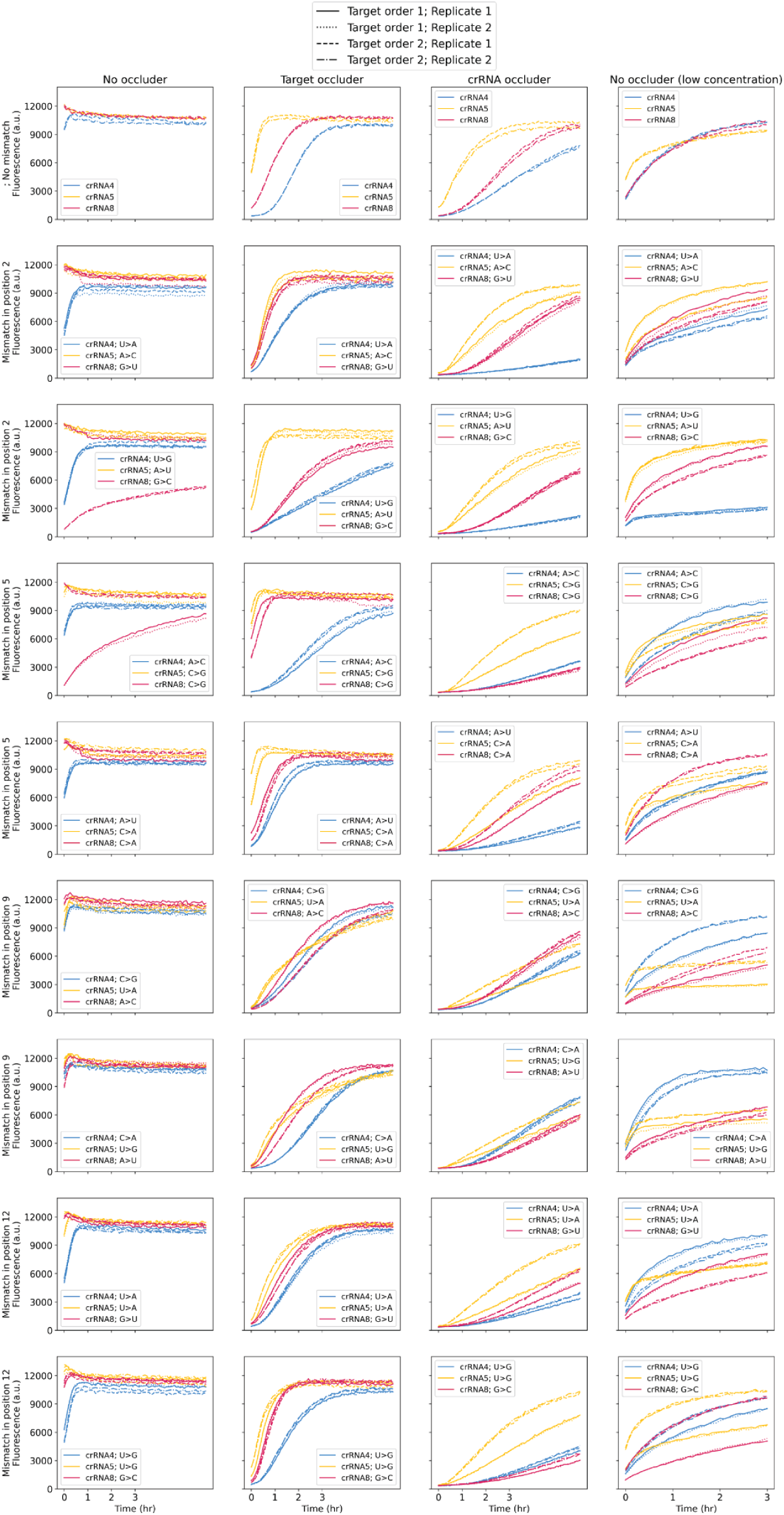
Mismatch detection using different types of occlusion. Raw fluorescence curves corresponding to Fig. 4A-B detecting various mismatches within three different target sequences for no occlusion, target occlusion, crRNA occlusion, and target+crRNA occlusion conditions.

**Extended Data Fig. 7:**
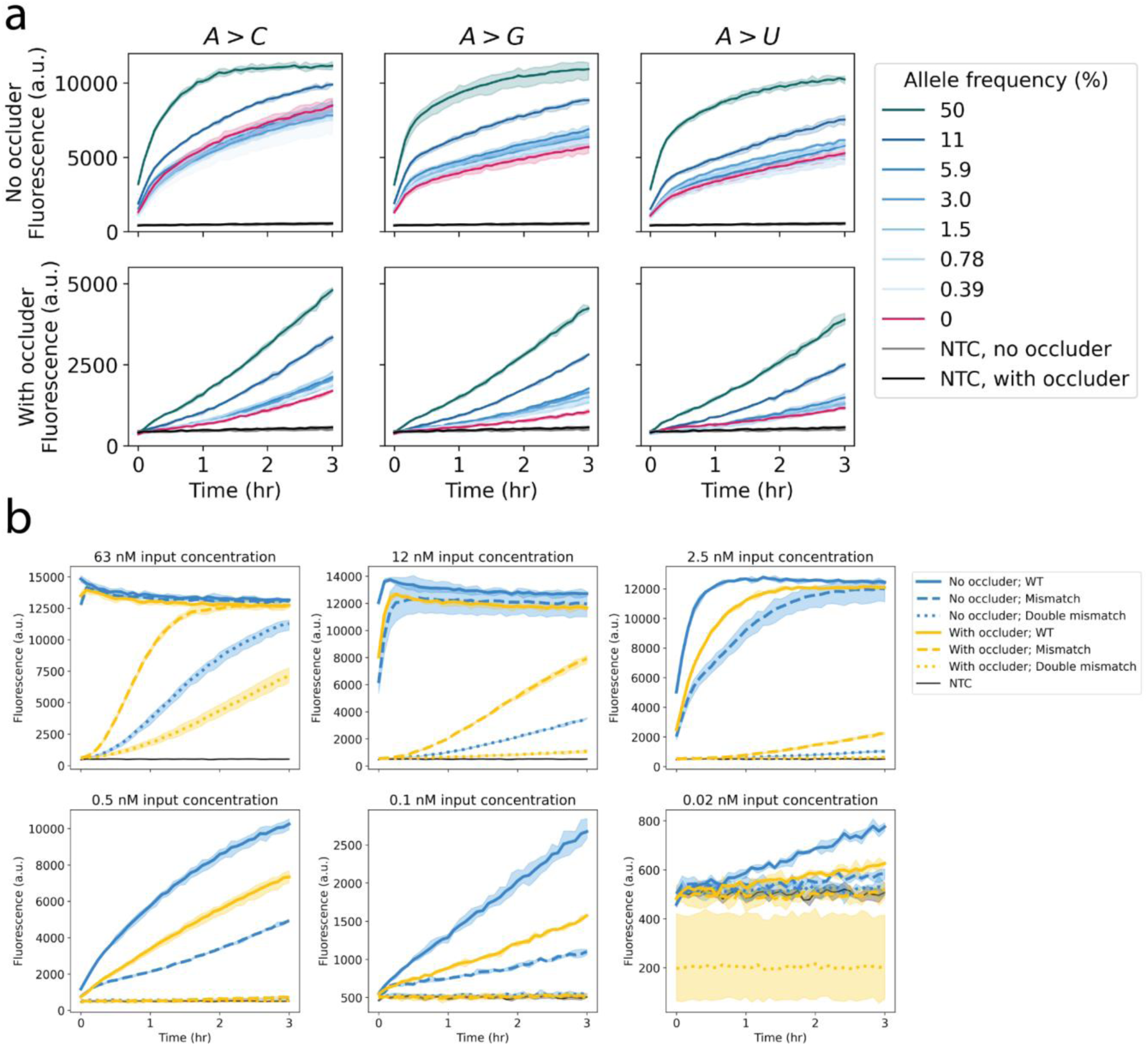
Fluorescence curves from experiments probing Cas13 mismatch detection at low allele frequencies and varying target concentration with occlusion. A. Raw fluorescence curves corresponding to Fig. 4C detecting mismatch detection sensitivity at decreasing allele frequency with and without occlusion. B. Raw fluorescence curves corresponding to Fig. 3D detecting presence of a mismatch at different target RNA concentrations with and without occlusion. Shaded regions show the range of fluorescence measurements for each condition across replicates.

**Extended Data Fig. 8:**
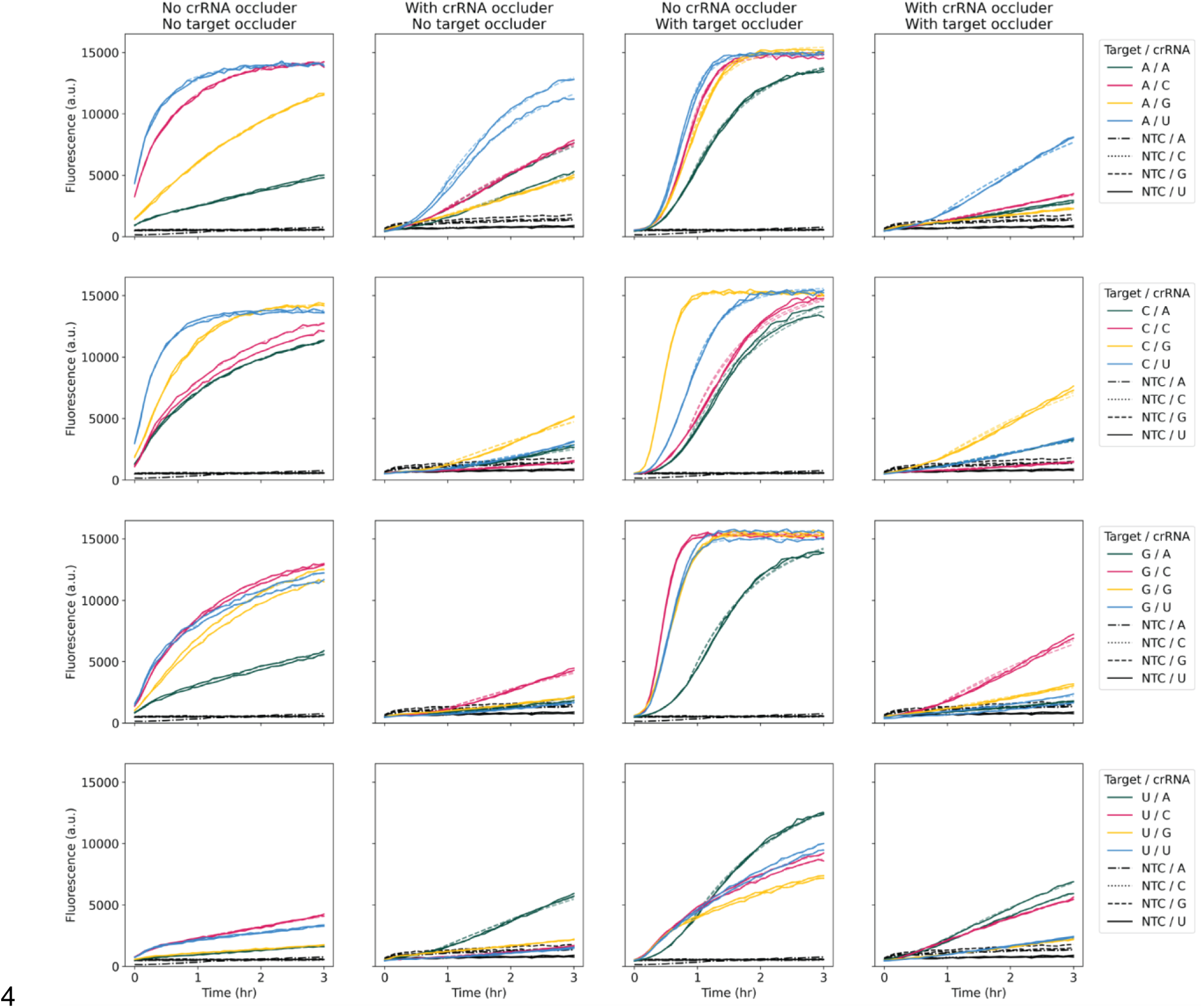
Fluorescence curves from Cas13 specificity matrix experiment with different types of occlusion. Raw fluorescence curves corresponding to Fig. 4D using all possible crRNA and target nucleotides at a given position. Semi-transparent dashed lines show curve fits (see Methods).

**Extended Data Fig. 9:**
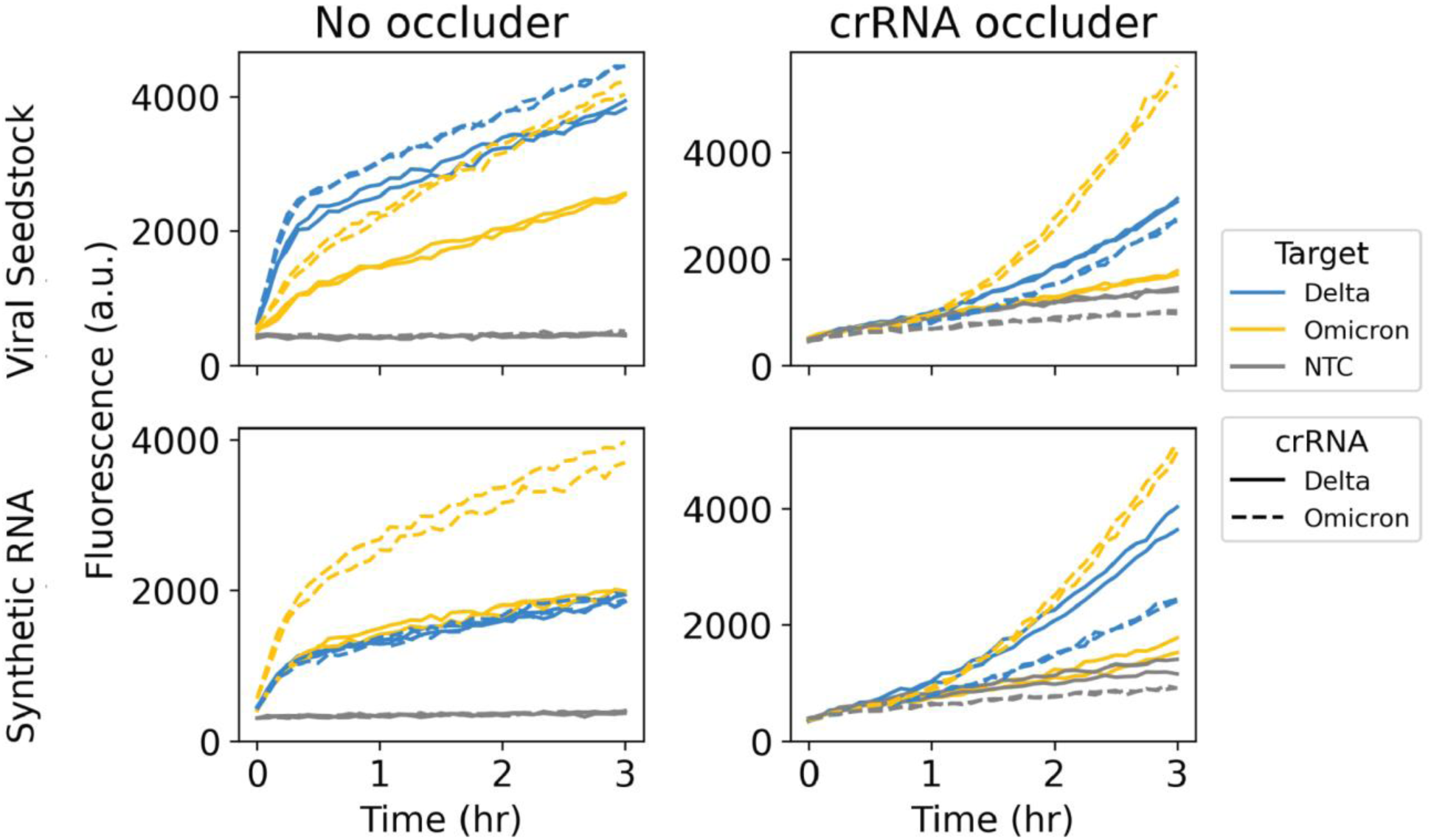
Fluorescence curves for SARS-CoV-2 variant calling experiments. Raw fluorescence curves corresponding to Figs. 4E and F targeting both Delta and Omicron variants with guides specific to each variant.

